# Modeling Organoid Population Electrophysiology Dynamics

**DOI:** 10.1101/2025.03.02.641081

**Authors:** Matthew J. Roos, Diego Luna, Dowlette-Mary Alam El Din, Thomas Hartung, Lena Smirnova, Alex Proescher, Erik C. Johnson

## Abstract

Improving models to investigate neurodegenerative disease, neurodevelopmental disease, neurotoxicology and neuropharmacology is critical to improve our basic understanding of the human nervous system, as well as to accelerate discovery of interventions and drugs. Improved models of the human central nervous system could enable critical discoveries related to functional changes induced by sensory stimulation or toxic exposures. Neural organoids, complex three-dimensional cell cultures derived from adult human stem cells, have been grown with complex connectivity and neuroanatomy. Moreover, these cultures have been interfaced with bi-directional electrical stimulation and recording, as well as chemical stimulation. This effort sought to develop new computational techniques which could be applied to comparative studies using neural organoids. In particular, we adapted CEBRA, a state of the art model from the *in vivo* modeling literature, to generate 2D and 3D embeddings (projections into structured low-dimensional spaces) of high dimensional neural organoid electrophysiology data. This can be done in an unsupervised or semi-supervised manner. Results indicate these embeddings can be quickly and reliably generated and serve as a low-dimensional, interpretable embeddings for characterizing changes in neural organoid activity over time, as well as clustering results around known bursting phenomena. Moreover, we demonstrate that mixtures of von Mises-Fisher distributions can be used as a parametric model for these embedding spaces to enable statistics hypothesis testing. This technique may enable new types of comparative studies using neural organoids, and may be critical for creating a representation for quantitative comparison and validation of neural organoid models against human and animal data. Looking ahead, this work could allow the formulation of a new class of experiments investigating the functional impact of toxins, genetic manipulations, or pharmacological interventions on human neurons.

## 1 Introduction

The ability to artificially replicate the function of the human brain in the laboratory, or *in vitro*, is a grand challenge for modern biology. Towards this goal, emerging research has made progress towards creating model systems that substantively aid our understanding of neurodevelopmental and neurodegenerative disorders and the interaction of chemical and biological threats with the nervous system. These systems, often dubbed microphysiological systems or neural organoids (1), are cultured from adult human induced Pluripotent Stem Cells (iPSCs). Bioengineering and stem cell technologies have led to the creation of a wide range of neural organoid systems capable of replicating the cell types of cortex, hippocampus, and more (2; 3; 4; 5), are primarily used in pharmacology and toxicology for high-throughput testing. Repeatable protocols have enabled development of large numbers of neural organoids for testing (6). As neural organoids grow in size and neuroanatomical fidelity, a key question remains: how similar is the activity of neural organoids to *in vivo* systems? This is a key prerequisite for using neural organoids as a model to explore aspects of functional processing.

It is well known that while even individual neurons have a range of complex and dynamic functions (7), at a larger scale, novel population dynamics emerge, such as brain rhythms (8). There are many mechanisms which may give rise to stable brain rhythms and complex population dynamics, such as specialized connectivity structures, and the role of microglia, astrocytes and oligodendrocytes in homeostasis (9). Progress in creating larger neural organoids (towards the size of a mouse brain) has resulted in increased complexity in terms of composition and activity. With that complexity comes the need to be able to detect and monitor nuanced changes in the activity that arises from chemical and electrical stimulation. In the context of neurotoxicity or neuropharmocology, changes from normal to abnormal physiological behavior may be subtle, and require sophisticated methods to model and predict such behavior. Moreover, a critical capability required is the ability to qualitatively and quantitatively compare the responses of neural organoids to *in vivo* systems, which may be recorded with very different sensors and tested with different experimental paradigms.

Electrophysiology, recording the electric biopotentials of neurons, remains one of the key methods for interrogating brains with the highest temporal and spatial resolution. While many studies utilize genomics and microscopy to study neural organoids, electrophysiology is a preferred method for studying neural population activity at the millisecond time scale. Many systems exist for *in vivo* and *in vitro* electrophysiology with various morphologies, channel counts, and impedances. Neural organoid cultures are often recorded with planar 2D microelectrode arrays (MEAs) (10) suitable for *in vitro* cultures only. In comparison, many *in vivo* studies are conducted with probe electrodes inserted into the brain, such as high-density Neuropixels (11). Novel electrode morphologies, such as 3D flexible electrodes (12), are rapidly emerging as tailored approaches for neural organoids. Comparing across these systems has enabled preliminary insight, for instance the relation of organoid burst frequencies to frequencies of neural rhythms during early development (13). However, in general, these electrophysiological recordings have different physical parameters, electrical properties, channel counts, and are applied to very different physiological systems. Arbitrary raw recordings cannot necessarily be used together for meaningful comparative analysis.

Given the rapid growth in high channel-count and high throughput methods for acquiring *in vivo* electrophsyiology, there has been considerable progress in approaches to model and extract insight from these highly stochastic, high dimensional recordings. These approaches aim to study different brain regions and the encoding of key behavioral variables (position, sensory information) in neural populations. For example, an experiment may record from hippocampus during a navigation task, aiming to study how hippocampal populations represent spatial positions. While traditional unsupervised dimensionality reductions approaches such as principle component analysis, independent component analysis, or non-negative matrix factorization (14) could be applied, they fail to capture the complex spatial and temporal dynamics in real data. New approaches are emerging which utilize deep learning models to create feature vectors corresponding to windows of time. These approaches are typically trained in a self-supervised fashion using a contrastive learning loss, which penalizes nearby embeddings of distinct time windows (e.g., windows separated by a great deal of time) (15; 16). These approaches are being used for neural latent modeling: estimating the latent variables (position, state, intent) encoded in neural populations. A new machine learning community has emerged to study these *in vivo* recordings, including the creation of the Neural Latents Benchmark (17). Of particular note is the CEBRA approach (16), a contrastive learning tool for training neural networks to create low-dimensional embedding spaces, which can be used in a fully unsupervised manner (utilizing only time) or with behavioral labels relevant to experiments. This approach is showing considerable success in creating interpretable representations of large, high-dimensional *in vivo* recordings.

There has been considerable interest, but less progress, in developing similar computational approaches to understand the behavior of neural organoids. It is envisioned that computational approaches could be used to develop predictive models of neural organoid activity, analysis tools for experimental results, and ultimately computational surrogates for *in silico* experimentation. These approaches may result in the creation of New Approach Methodologies (NAMs) (18; 19; 20) for neurotoxicity and neurodevelopmental studies utilizing neural organoids. Such approaches may also form the basis for “digital twins” of neural organoids, as have been developed for other types of *in vitro* systems (21). There is considerable interest in developing computational approaches to understand and predict the dynamics of complex neural organoid physiology.

In this effort, we aimed to address the lack of computational approaches to study neural organoids which incorporate the latest insights from deep learning methods. Latent space embedding approaches, such as CEBRA, have not been widely explored for neural organoids. Driven by the need to characterize and compare *in vivo* and *in vitro* systems, we curated a dataset and extended the CEBRA approach to neural organoids, creating MOPED (Modeling Organoid Population Electrophysiology Dynamics). In this work, for the first time, we utilized a contrastive latent space model (CEBRA) to train neural organoid embedding models. We used these to characterize trajectories within a recording session, across different ages of neural organoids, and across days of recording. Additionally, we applied parametric statistical models to create parameterized fits suitable for hypothesis testing. We believe this technique will be critical to explore experimental paradigms for neural organoids to enable new studies into neurodevelopmental disorders, neurodegenerative disease, neural toxicology, and neural pharmacology.

## 2 Methods and Models

To develop and test the MOPED framework, we built closely on the CEBRA methodology and incorporated several neural organoid and animal electrophysiology datasets into a common framework (Fig. 1).

**Figure 1:**
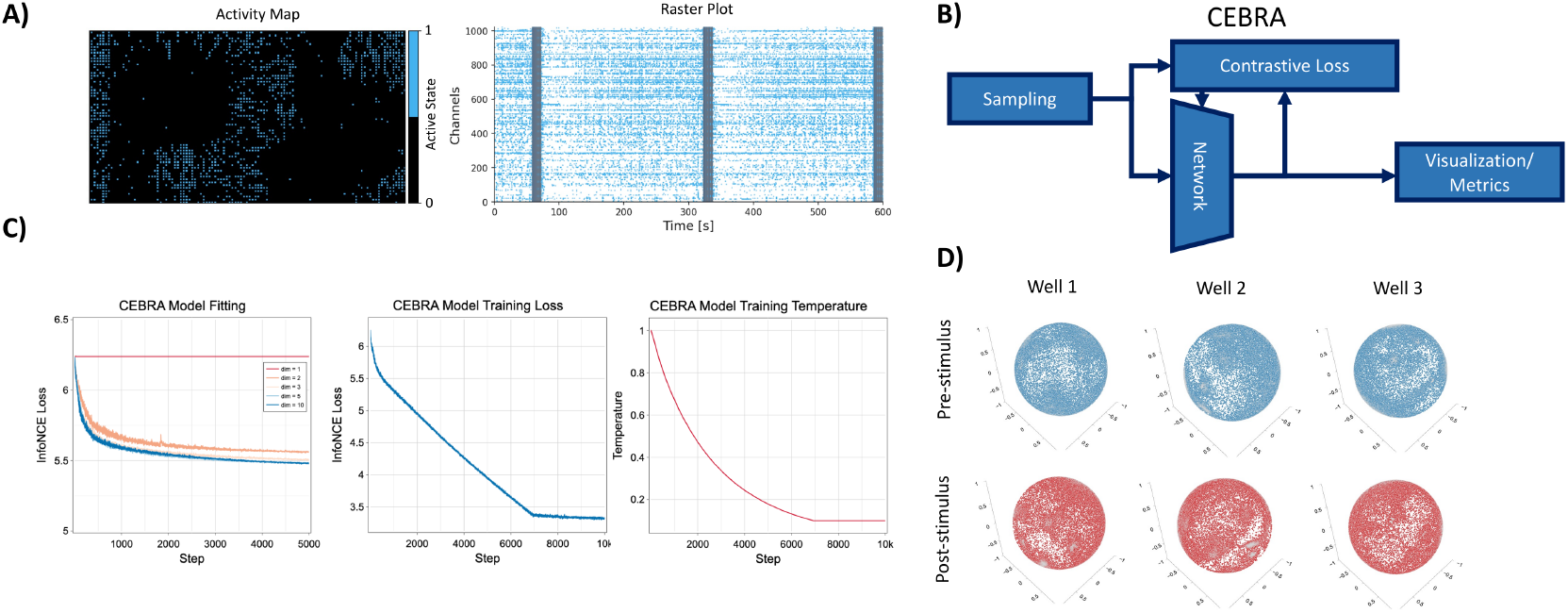
Overview of the MOPED system, incorporating the CEBRA framework, for neural organoid data. Panel A) shows examples of electrophysiology recordings utilizing a 2D MEA system. This shows a 2D representation of the active electrodes on the array, and an example spike-time raster plot. Panel B) gives an overview of the MOPED framework, where the MOPED dataloader produces batches for the CEBRA network, which is optimized to minimize a contrastive loss function. Panel C) shows example curves for training the CEBRA network, showing loss for different numbers of dimensions, as well as the adjustment of training temperature. Of note is the smooth convergence of the CEBRA model for neural organoid data when using only time labels. Panel D) shows example 3D embeddings generated from three wells pre- and post-stimulus. The distribution shifts indicate possible changes in functional activity due to stimulation, which can also be investigated using this method.

### 2.1 Neural Organoid Electrophysiology

In this effort, we curated a set of neural organoid electrophysiological recordings which were collected at the JHU Bloomberg School of Public Health to investigate mechanisms of learning and memory (22).

Briefly, female fibroblast-derived N8 iPSCs were obtained from the National Institute for Biological Standards and Control. These cells were cultured in mTESR-Plus medium (StemCell Technologies). Cells were then differentiated in a monolayer to neuroprogenitor cells using a serum-free, neural induction medium (Gibco, Thermo Fisher Scientific). Nestin/Sox2-positive cells were then expanded and seeded in uncoated 6-well plates. These cultures were incubated and subject to constant gyratory shaking to form spheres.

Data in this study were taken from two or three independent cultures with different ages, and included a “time-course experiment” of recording spontaneous activity over a three week period and a “stimulation” dataset consisting of theta burst stimulation of neural organoids (23).

Organoids were plated on six-well plates for an integrated MEA amplifier (Maxwell Biosystems, Zurich, Switzerland). Two groups of organoids were plated-one group at age 40-44 days, and one group at age 68-70 days. The age group plated at age 40-44 days has two independent groups (differentiations of organoids). One group had 5 wells and one group had 6 wells (for a total of 11 wells for this age group). The age group plated at 68-70 days has three independent groups (differentiations of organoids). One with 6 wells, another with 6 wells, and the last with 12 wells (for a total of 24 wells for this age group). For the time-course experiment, organoids were recorded in 10 minute sessions per day, for three weeks. For the stimulation experiment, organoids were recorded for a baseline 10 minute period, stimulated, then recorded for four hours post-stimulation. Up to 1024 channels were recorded at 10kHz, which were saved as hdf5 files for further secondary analysis.

### 2.2 *In Vivo* Electrophysiology

In addition to the neural organoid electrophysiolgy dataset, this effort curated a range of open-source animal electrophysiology datasets which can be used for pretraining and comparative studies. These included datasets from rat and macaque studies. These datasets are publicly available on the DANDI archive, formatted using the Neurodata Without Borders (NWB) format (24). For this study, a set of recordings from rat hippocampus during the sleep/wake cycle (25) were used as an *in vivo* comparison dataset.

For future work, several datasets utilized in the Neural Latents Benchmark (17) could be valuable for comparison between organoid and mammalian models using the CEBRA technique. These datasets include recordings from rat hippocampus during navigation (26), recordings during non-human primate reaching tasks (27; 28), and recordings during Macaque interval reproduction task (29).

These datasets provide validation references for methodological improvements, pre-training datasets for neural organoid modeling, and comparative datasets to explore the link between *in vitro* and *in vivo* models.

### 2.3 MOPED Dataset Organization and Data Loaders

Given these curated datasets, the MOPED effort created a Python framework for flexibly accessing these data for model training and computational experiments. Data were provided in NWB and hdf5 formats with different metadata. The MOPED effort created a JSON-based dataset specification which can be used to capture the structure of dataset files as well as corresponding metadata related to samples, recording, and experimental conditions. This framework can be flexibly extended to create new standards which incorporate in additional fields to support new experiments. From this, an experiment-specific YAML file can be specified, which indicates how files should be sampled for a given experiment, including train and test splits and other limits on valid time lengths and channel counts. These configuration files, along with the raw data files, are read by the MOPED Pytorch data loader, which creates batches for machine learning training. These batches are then input into the CEBRA framework for training and inference. For both *in vivo* and *in vitro* networks, the inputs are temporal windows of spike-time data, derived from the spike sorting and spike detection methods applied to various datasets.

### 2.4 CEBRA for Neural Organoids

The MOPED framework builds upon the CEBRA approach for learning embedding spaces (16), which has been widely employed for modelling neural data.

Briefly, two data samples *x* and *y* are mapped into an embedding space of fixed dimensionality *D* (typically 2D or 3D) through neural networks *f* and *f* ^′^. The neural networks are trained to minimize the expected loss given a similarity measure *ϕ* and temperature parameter *τ* (which is reduced over epochs during training). The loss function, minimized in expectation over a sequence of length *n* of samples of *x, y* is

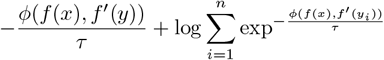

This can be minimized using standard approaches for gradient descent. Embedding functions are normalized to create a distribution on the unit hypersphere of dimension *D*, and temporal sequences can be plotted for sequences of inference samples *x*_*t*_.

The key issue in adapting CEBRA to neural organoid data is deciding on approaches for sampling pairs *x, y*. In this work, we utilize the simplest and most general approach, sampling only using temporal information (short versus long intervals between samples). A critical future pathway is exploring approaches to utilize experimental information (age, condition) and cross-dataset information (species, recording method) to improve the insights provided by the embedding approach.

### 2.5 Metrics and Analysis

In addition to new dataloader and data management techniques, the MOPED framework also adds additional metrics and analysis to the study of embedding spaces created by CEBRA. These are focused on providing quantitative tools to compare neural organoid datasets with each other, as well as *in vivo* data. First, we compute basic electrophysiology statistics including spike count and burst statistics. Bursts were detected and characterized using a variant of the MaxInterval method with hand-tuned thresholds (30).

In addition, embedding space distributions were characterized in three key ways. First, the entropy of the distribution (− Σ*p*(*x*) log *p*(*x*)) was computed. When labels are available for particular time points (in this case derived from the burst detection algorithm) the silhouette score can be computed to assess the degree of clustering for different conditions. Finally, we fit a mixture of von Mises-Fisher distributions (31), a parametric distribution over the unit hypersphere. Different numbers of components can be specified, and these fits can be used for parametric hypothesis testing and likelihood ratio calculations.

## 3 Results

The MOPED CEBRA framework was used to train embedding models for *in vivo*, including example recordings from the sleep-wake cycle recorded from rat hippocampus. In addition, dozens of neural organoid recordings were processed to create 2D and 3D embeddings of data across days *in vitro*, as well as pre- and post-stimulus.

### 3.1 Fitting CEBRA Models to Neural Organoid Data

To begin exploring the MOPED method and training CEBRA models for neural organoids, we conducted several test fits of data pre- and post-stimulus from the stimulation dataset (Fig. 1). This shows the neural organoid electrophysiology data, represented as a map of active electrodes on the 2D MEA, as well as spike-time rasters of the detected spike events over the 1024 channels. The complexity of the high-dimensional data can be seen, as well as the presence of lower-dimensional burst events (indicated by the strong vertical lines of activity in the spike raster). CEBRA models could be consistently fit for a wide range of neural organoid datasets and hyperparameters, showing well-behaved loss curves (Fig. 1C). After training, the training sample or new samples from a test dataset can be visualized as a 2D or 3D embedding to study the properties learned by the embedding model. Figure 1D shows three example wells pre- and post-stimulus, indicating a change in the embedding distribution between the two conditions. The MOPED approach can be used to train and create these latent space embeddings from a wide range of experimental data.

### 3.2 Characterization of Neural Organoid Embeddings Across Days

To begin investigating the intepretability and utility of these latent spaces, we generated latent space embeddings for each recording in the time-course dataset (consisting of two populations of neural organoids, of different ages, recorded over three weeks). These populations both showed shifts in spike rate over time, following a roughly linear trajectory, but with a difference in slope between the two age groups (Fig 2A). Applying the MOPED approach (Fig. 2B and c), latent space embeddings were created independently for each well and day for the two populations. From these, the entropy of the embedding distribution was calculated. The entropy of the embedding distribution increases over time for both populations in a manner similar to spike rate increases. However, the younger organoids show a clear saturation in the entropy of the embedding space distribution after several days despite continually increasing spike rates. This suggests both key correlations and differences between the changes in the embedding space distribution and basic spike statistics over time.

**Figure 2:**
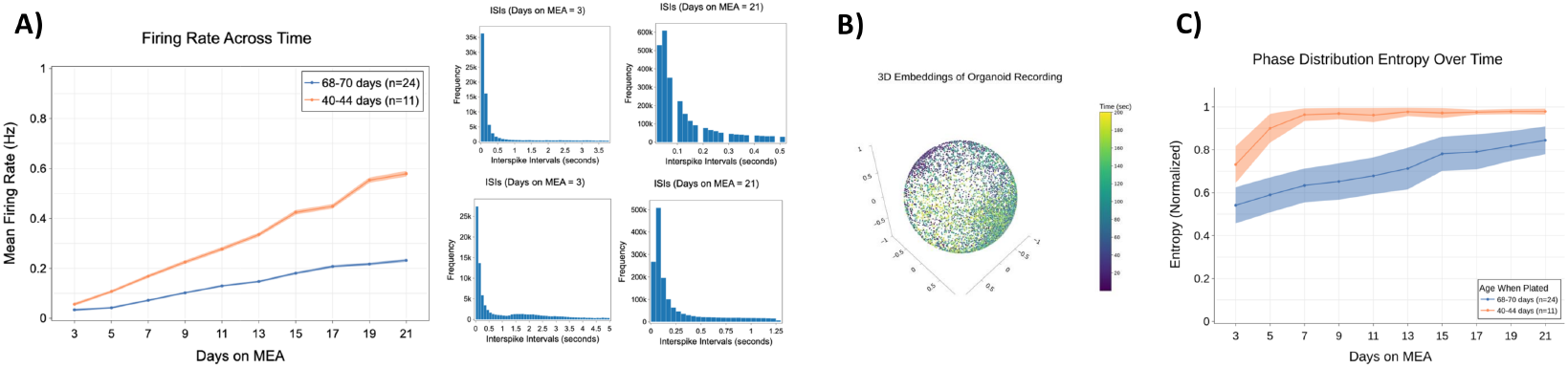
Characterization of neural organoid spike statistics and entropy of 3D neural organoid embeddings spaces. Panel A) characterizes the changes in spike firing rate estimated across six wells for two separate groups (6 wells each) of older and younger neural organoids. Recordings took place over 3 weeks. There is a clear linear trend in the spike rate for both groups. Also shown are the ISI distributions for an example well over four days. Panel B) shows an example latent space embedding calculated from an organoid plated at 70 days and recorded at day 13. Color coding shows the trajectories of activity over a 200 second period. Panel C) shows the entropy for the distribution of samples in the 3D latent space, calculated for the group of younger and older organoids plotted over the same three week period as Panel A). We can see an increase in entropy, which saturates for the younger organoids early in the recording period.

### 3.3 Characterization of Neural Organoid Burst Activity

To investigate the ability of the MOPED embedding approach to characterize key states in neural organoid data, we investigated a known phenomena in neural organoid recordings, bursting (Fig. 3). Bursts are well known to occur in these cultures, and burst statistics change over time (Fig. 3B). In an unsupervised fashion, 3D embedding spaces were trained for these recordings using the MOPED approach. Plotting a single embedding dimension versus time, there is a clear correlation of peaks in the embedding value with burst activity, which is maintained in the test dataset (Fig. 3A). In the embedding space, time periods corresponding to bursts (detected using the burst detector with hand-tuned thresholds), roughly cluster into one portion of the embedding space (Fig. 3C). Computing the silhouette score, it can be seen that the bursts are reasonably well clustered (a score of -1 indicates a uniform, unclustered distribution). The technique shows a difference in burst clustering between the two age datasets, and the silhouette score is roughly constant over time, following the trend of burst duration and burst interval. Overall, this result suggests the MOPED approach can be used in an unsupervised manner to discover clusters related to key features in the neural organoid dataset.

**Figure 3:**
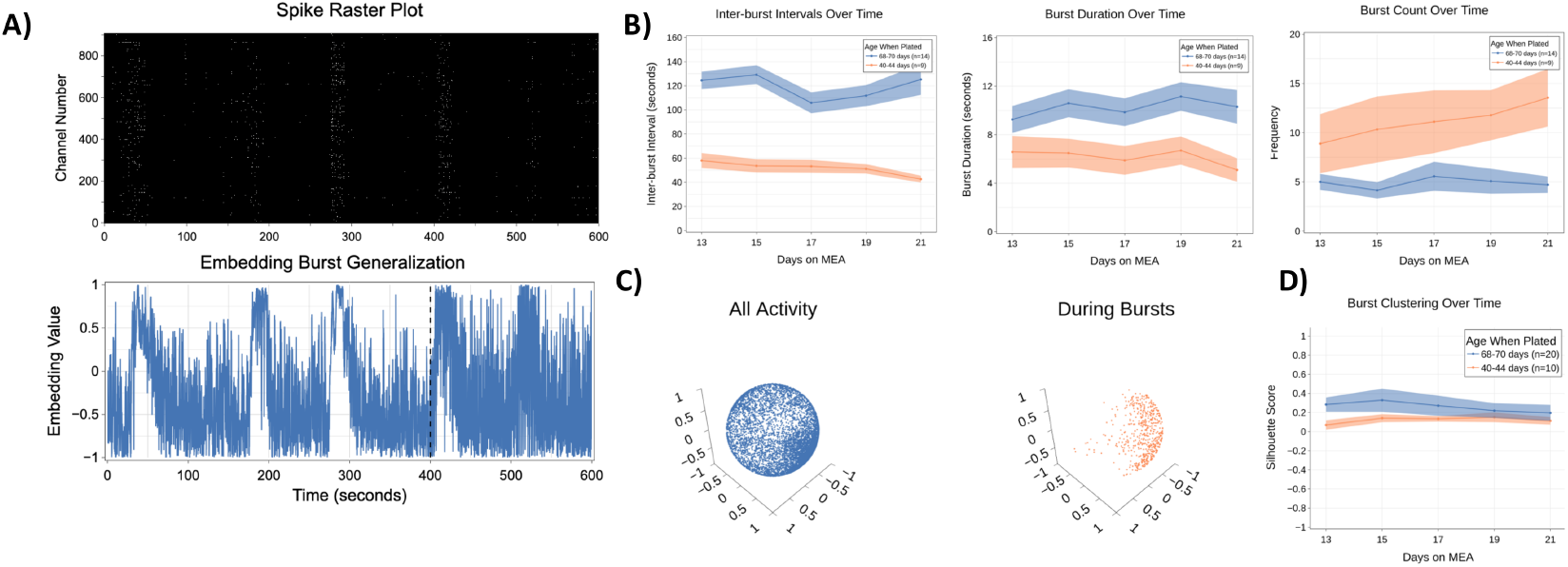
Characterizing burst events using the MOPED approach as an unsupervised data analysis tool. Panel A) shows an example time sequence from a neural organoid demonstrating bursts in the spike raster plot. A corresponding embedding vector (learned in an unsupervised fashion) for one dimension is plotted over time, where peaks correspond to bursts. The dashed line indicates the train-test split for this recording, and it can be seen that peaks continue to correspond to bursts in the test dataset. Panel B) shows the burst statistics derived from a hand-tuned, threshold-based burst detector. Recordings with fewer than 4 bursts have been filtered out before computing burst statistics, leaving 14 and 9 valid recordings from each group. This shows general consistency in the rate of bursts, with a distinct difference between younger and older organoids. Panel C) shows an example 3D embedding derived using the MOPED approach, including a color coding of time points corresponding to the output of the burst detector, indicating clustering. Panel D) shows the silhouette score for the clusters using all recordings (using the ground truth labels) over days, indicating a difference between the young and old populations and demonstrating considerable, though not perfect, clustering.

### 3.4 Parametric Fits of Embedding Distributions

In addition to generating latent space embeddings, qualitatively visualizing, and computing entropy, fitting parametric distributions can be a powerful tool for quantitative, comparative analysis. In particular, these approaches enable parametric statistical tests, improved calculation of likelihood ratios, and intepretable comparison of distributions. The mixture of von Mises-Fisher distributions is an appropriate choice as it is formulated for unit hyperspheres and allows for specification of the number of components for modeling (similar to Gaussian mixture models). Figure 4 shows examples for 2D and 3D fits of neural organoid embeddings. The fits show that the distribution can qualitatively capture the features of the data histogram, and that increasing the component count improves the fit of the data. As usual with such techniques, model selection criteria (32) should be applied to balance model complexity with model goodness-of-fit.

**Figure 4:**
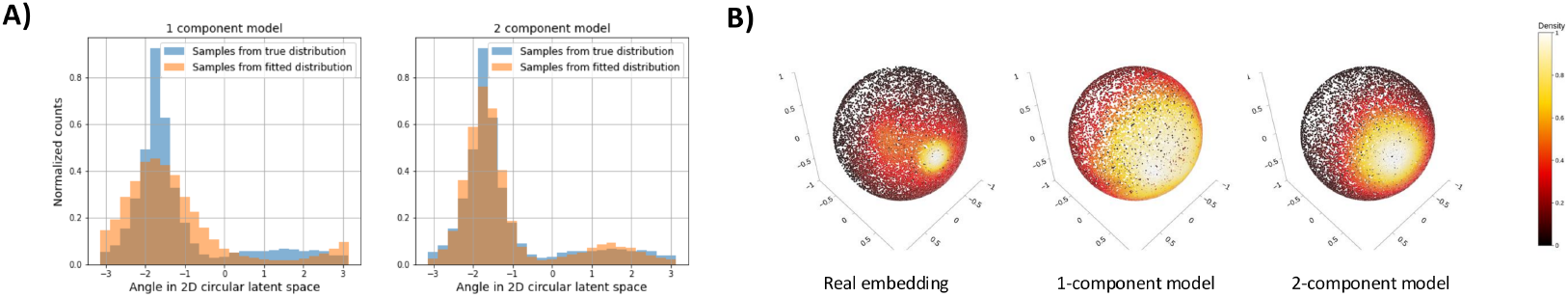
Demonstration of the use of von Mises-Fisher distributions to fit parametric models for neural organoid latent spaces. Panel A) demonstrates 2D fits to data from and example well, showing the fit for a single component and two component model. Panel B) demonstrates similar fits for 3D data from the same exemplar well, for a one and two component model.

### 3.5 Comparison of Neural Organoid and *In Vivo* Embeddings

One of the advantages of the MOPED approach, enabled by the underlying CEBRA model, is that low-dimensional embedding spaces can be learned from data with very different underlying statistics, channel counts, and recording methodologies. As a preliminary demonstration of this, we show 2D embeddings of wake-sleep data from rat hippocampus learned through the MOPED framework (Fig. 5). These indicate distinct embedding distributions for the sleep wake data, and measures of entropy, as well as von-Mises Fischer distribution fits, can be calculated. This allows direct comparison to similar embeddings from neural organoids (Fig. 4). However, it is critical to recognize that the generated embeddings from CEBRA may not be aligned between datasets. Therefore a fitting approach should first be applied to compute the rotation that maximizes alignment before fitting the mixture of von-Mises Fisher distributions. While preliminary, this suggests a path towards comparative analysis of *in vitro* and *in vivo* data using the MOPED framework.

**Figure 5:**
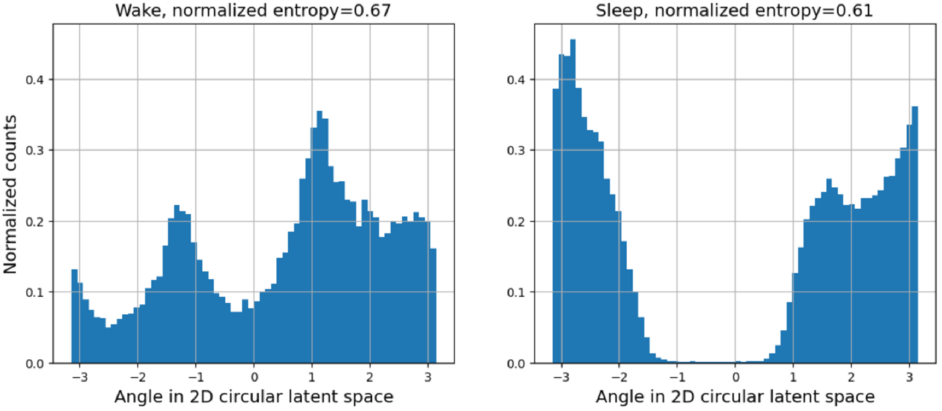
Example 2D embeddings of *in vivo* data derived from rat hippocampal recordings in a dataset including sleep and wake states. The left panel shows the distribution for the Wake state, while the right panel shows the distribution for the Sleep state. There are clear differences in the shape and peaks of the distributions, as well as the uniformity of the distributions. Despite having different numbers of electrodes (64 electrodes used *in vivo* versus 1024 used *in vitro*) and different recording parameters, a comparable 2D representation is created.

## 4 Discussion

The MOPED framework presented here is a flexible and general approach to leverage CEBRA to model neural organoids and *in vivo* electrophysiology datasets. The framework allows for extensible specification of datasets and experiments in text files suitable for version control to encourage experimental reproducibility. The framework can be trained for large numbers of recordings, consistently converging to solutions which generalize from train to test time windows. While additional efforts are required to tune hyperparameters and design experiments for neural organoids, training complexity is suitable for workstations or single compute nodes. It is anticipated in the coming years this will rapidly grow into hundreds of terabytes of data, which will greatly benefit from rapid tools for machine learning analysis.

The MOPED framework has several key advantages compared to previous analysis approaches. Traditional measures of spike-time statistics and burst detection, can provide insight but are limited in their ability to capture complex time-varying phenomena. Moreover, deriving insight from these analysis often take considerable manual analysis, especially when systems and conditions vary considerably over time and experimental conditions. Data-driven approaches, such as principle component analysis, independent component analysis, non-negative matrix factorization, and similar approaches, can be applied to these data. However, many of these approaches have significant assumptions of the statistics or structure of components, the neural network embedding approach of CEBRA generalizes this. Moreover, the contrastive learning scheme allows for incorporation of behavioral or experimental variables which cannot be included in these traditional analysis frameworks.

One of the most promising aspects of the MOPED (and CEBRA) approach is the ability to generate interpretable comparisons between different systems and experiments, which may vary widely in channel count, spike statistics, coverage, and other key parameters. While these systems may have trivially different basic statistics (spike rate, channel count, etc), the key issue of experimental interest is if behavior or task-related variables are encoded, and how these are modified by experimental conditions or treatments. The low-dimensional latent spaces generated by MOPED may provide a key approach for comparative studies seeking to gain insight from neural organoids, and ultimately augment or replace human and animal studies. Key innovations introduced in the MOPED approach are quantitative metrics of entropy as well as the parametric fit of a mixture of von-Mises Fisher distributions. These approaches may allow for rapid comparative experiments, including hypothesis testing.

Taken together, the MOPED approach can be combined with existing electrophysiological analysis techniques to enable new types of computational experiments using neural organoids (Fig. 6). In this envisioned framework, neural organoids exposed to different conditions and stressors can be generated for experimentation. Testing tasks, drawn from behavioral animal and human experiments, can be adapted to provide stimulation for the organoids. Electrophysiological recordings from the organoids, including single unit activity, local field potentials, and task performance, can be recorded for analysis. Optionally, comparative datasets of animal and human electrophysiology can be included, for both validation and comparative analysis. The MOPED analysis approach can join other analysis techniques, such as network analysis and plasticity analysis, to form a bank of functional activity assays for comparative studies. In the context of neural toxicology, neural pharmacology, and cognitive studies, this framework may provide a fundamentally new approach for deriving scientific insight compared to human and animal studies.

**Figure 6:**
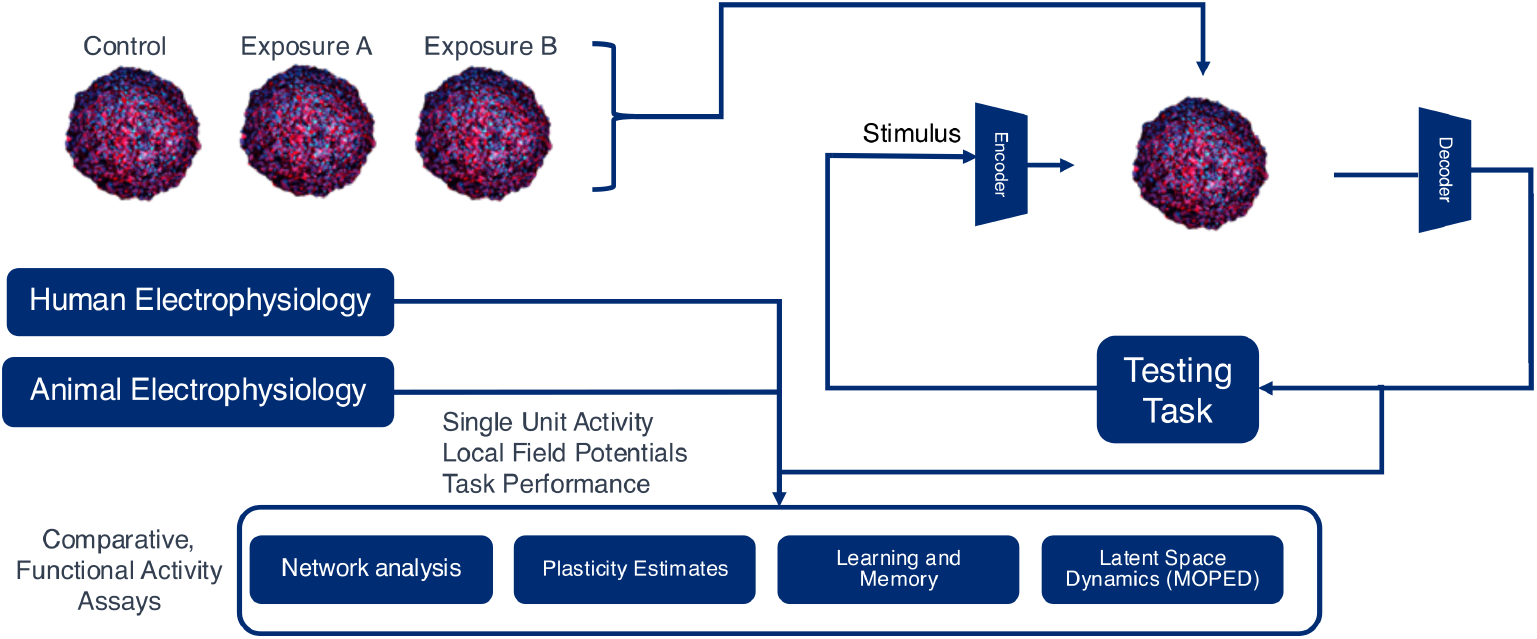
Envisioned future concept for human neural organoid experimentation. Control organoids and neural organoids exposed to conditions relevant to neurological health can be used in a testing framework which stimulates and records to elicit functional activity based on a test task. Data from the recordings can be combined with human and animal data, if desired, for validation or comparative studies. These data can then be assessed with a bank of functional activity assays including, but not limited to, the proposed MOPED technique. Experiments of this type may represent a compelling alternative to studying neurotoxicological and neuropharmacological effects.

## 5 Conclusions

The impact of this work, in conjunction with continuing advancements in the size and bio-fidelity of neural organoids, will be to substantially advance the ability of researchers to assess the neurotoxicity of chemical compounds to which people may be exposed (33). The MOPED approach can be rapidly applied to a range of neural organoid and *in vivo* recordings. This modelling approach, combined with statistical metrics and models applied to the generated embedding spaces, can increase our ability to characterize normal and abnormal activity in neural organoids. In future experiments, when neural organoids are exposed to chemicals in the lab, the MOPED technique can be used in conjunction with existing approaches to gain insight into functional changes in human neural tissue. Continued work will be needed to advance the foundational aspects of the modeling approach as well as apply the MOPED technique to aid in understanding of neural activity patterns in neural tissue.

## Author Contributions

MR, DL, and EJ contributed to conceptualization, data curation, methdology, software, and writing. DA, TH, and LS contributed to data curation, conceptualization, writing - review and editing, and methodology. AP contributed to conceptualization, supervision, validation, project administration, and writing - review and editing.

## Acknowledgments

We would like to thank the authors and maintainers of the CEBRA repository.

